# Context-Aware Amino Acid Embedding Advances Analysis of TCR-Epitope Interactions

**DOI:** 10.1101/2023.04.12.536635

**Authors:** Pengfei Zhang, Seojin Bang, Michael Cai, Heewook Lee

## Abstract

Accurate prediction of binding interaction between T cell receptors (TCRs) and host cells is fundamental to understanding the regulation of the adaptive immune system as well as to developing data-driven approaches for personalized immunotherapy. While several machine learning models have been developed for this prediction task, the question of how to specifically embed TCR sequences into numeric representations remains largely unexplored compared to protein sequences in general. Here, we investigate whether the embedding models designed for protein sequences, and the most widely used BLOSUM-based embedding techniques are suitable for TCR analysis. Additionally, we present our context-aware amino acid embedding models (`catELMo`) designed explicitly for TCR analysis and trained on 4M unlabeled TCR sequences with no supervision. We validate the effectiveness of `catELMo` in both supervised and unsupervised scenarios by stacking the simplest models on top of our learned embeddings. For the supervised task, we choose the binding affinity prediction problem of TCR and epitope sequences and demonstrate notably significant performance gains (up by at least 14% AUC) compared to existing embedding models as well as the state-of-the-art methods. Additionally, we also show that our learned embeddings reduce more than 93% annotation cost while achieving comparable results to the state-of-the-art methods. In TCR clustering task (unsupervised), `catELMo` identifies TCR clusters that are more homogeneous and complete about their binding epitopes. Altogether, our `catELMo` trained without any explicit supervision interprets TCR sequences better and negates the need for complex deep neural network architectures in downstream tasks.

## 1 Introduction

T cell receptors (TCRs) play critical roles in adaptive immune systems as they enable T cells to distinguish abnormal cells from healthy cells. TCRs carry this important function by binding to antigens presented by major histocompatibility complex (MHC) and recognizing whether the antigens are self or foreign [1]. It is widely accepted that the third complementarity-determining region (CDR3) of the TCRβ chain is the most important in determining its binding specificity to epitope—a part of an antigen [2, 3]. The advent of publicly available databases of TCR-epitope cognate pairs opened the door to computational methods to predict the binding affinity of a given pair of TCR and epitope sequences. Computational prediction of binding affinity is important as it can drastically reduce the cost and the time needed to narrow down a set of candidate TCR targets, thereby accelerating the development of personalized immunotherapy leading to vaccine development and cancer treatment [4, 5]. Computational prediction is challenging primarily due to 1) many-to-many binding characteristics [6] and 2) the limited amount of currently available data.

Despite the challenges, many deep neural networks have been leveraged to predict binding affinity between TCRs and epitopes [7, 8, 9, 10, 11, 12]. While each model has its own strengths and weaknesses, they all suffer from poor generalizability when applied to unseen epitopes, not present in the training data [7, 13]. In order to alleviate this, we focus mainly on embedding, as embedding an amino acid sequence into a numeric representation is the very first step needed to train and run a deep neural network. Furthermore, a ‘good’ embedding has been shown to boost downstream performance even with a few numbers of downstream samples [14, 15].

BLOSUM matrices [16] are widely used for representing amino acids into biological-related numeric vectors in TCR analysis [7, 9, 11, 17]. However, BLOSUM matrices are static embedding methods as they always map an amino acid to the same vector regardless of its context. For example, in static word embedding, the word “mouse” in phrases “a mouse in desperate search of cheese” and “to click, press and release the left mouse button” will be embedded as the same numeric representation even though it is used in different contexts. Similarly, the amino acid residue `G` appearing five times in a TCR*β* CDR3 sequence CASGGTGGANTGQLYF may play different roles in binding to antigens as each occurrence has a different position and neighboring residues. The loss of such contextual information from static embedding may inevitably compromise model performances [18, 19].

Recent successes of large language models [14, 15] have been prompting new research applying text embedding techniques to amino acid embedding. Large language models are generally trained on a large text corpus in a self-supervised manner where no labels are required [18, 20, 21]. A large number of (unlabeled) protein sequences has been available via high quality and manually curated databases such as UniProt [22]. With the latest development of targeted sequencing assays of TCR repertoire, a large number (unlabeled) of TCR sequences has also been accessible to the public via online databases such as ImmunoSEQ [23]. These databases have allowed researchers to develop large-scale amino acid embedding models that can be used for various downstream tasks. Asgari et al. [24] first utilized Word2vec [20] model with 3-mers of amino acids to learn embeddings of protein sequences. By considering a 3-mer amino acids as a word and a protein sequence as a sentence, they learn amino acid representations by predicting the context of a given target 3-mer in a large corpus of surrounding ones. Yang et al. [25] applied Doc2vec [21] models to protein sequences with different sizes of k-mers in a similar manner to Asgari et al. and showed better performance over sparse one-hot encoding. One-hot encoding produces static embeddings, like BLOSUM, which leads to the loss of positional and contextual information.

Later, SeqVec [26] and ProtTrans [27] experimented with dynamic protein sequence embeddings via multiple context-aware language models [18, 19, 28], showing advantages across multiple tasks. Note that the aforementioned amino acid embedding models were designed for protein sequence analysis. Although these models may have learned general representations of protein sequences, it does not necessarily signify their generalization performance on TCR-related downstream tasks.

Here, we explore strategies to develop amino acid embedding models and emphasize the importance of using “good” amino acid embeddings for a significant performance gain in TCR-related downstream tasks. It includes neural network depth, architecture, types and numbers of training samples, and parameter initialization. Based on our experimental observation, we propose `catELMo`, whose architecture is adapted from ELMo (Embeddings from Language Models [18]), a bi-directional context-aware language model. `catELMo` is trained on more than four million TCR sequences collected from ImmunoSEQ [23] in an unsupervised manner, by contextualizing amino acid inputs and predicting the next amino acid token. We compare its performance with state-of-the-art amino acid embedding methods on two TCR-related downstream tasks. In TCR-epitope binding affinity prediction application, `catELMo` significantly outperforms the state-of-the-art method by at least 14% (absolute improvement) of AUCs. We also show `catELMo` achieves an equivalent performance to the state-of-the-art method while dramatically reducing downstream training sample annotation cost (more than 93% absolute reduction). In the epitope-specific TCR clustering application, `catELMo` also achieves comparable to or better cluster results than state-of-the-art methods.

## 2 Results

`catELMo` is a bi-directional amino acid embedding model that learns contextualized amino acid representations (Fig. 1a), treating an amino acid as a word and a sequence as a sentence. It learns patterns of amino acid sequences with its self-supervision signal, by predicting each the next amino acid token given its previous tokens. It has been trained on 4,173,895 TCR*β* CDR3 sequences (52 million of amino acid tokens) from ImmunoSEQ [23] (Table 1). `catELMo` yields a real-valued representation vector for a sequence of amino acids, which can be used as input features of various downstream tasks. We evaluated `catELMo` on two different TCR-related downstream tasks, and compared its performance with existing amino acid embedding methods, namely BLOSUM62 [16], Yang et al. [25], ProtBert [27], SeqVec [26], and TCRBert [29]. We also investigated various components of `catELMo` in order to account for its high performance, including the neural network architecture, layer depth and size, types of training data, and the size of downstream training data.

**Figure 1.**
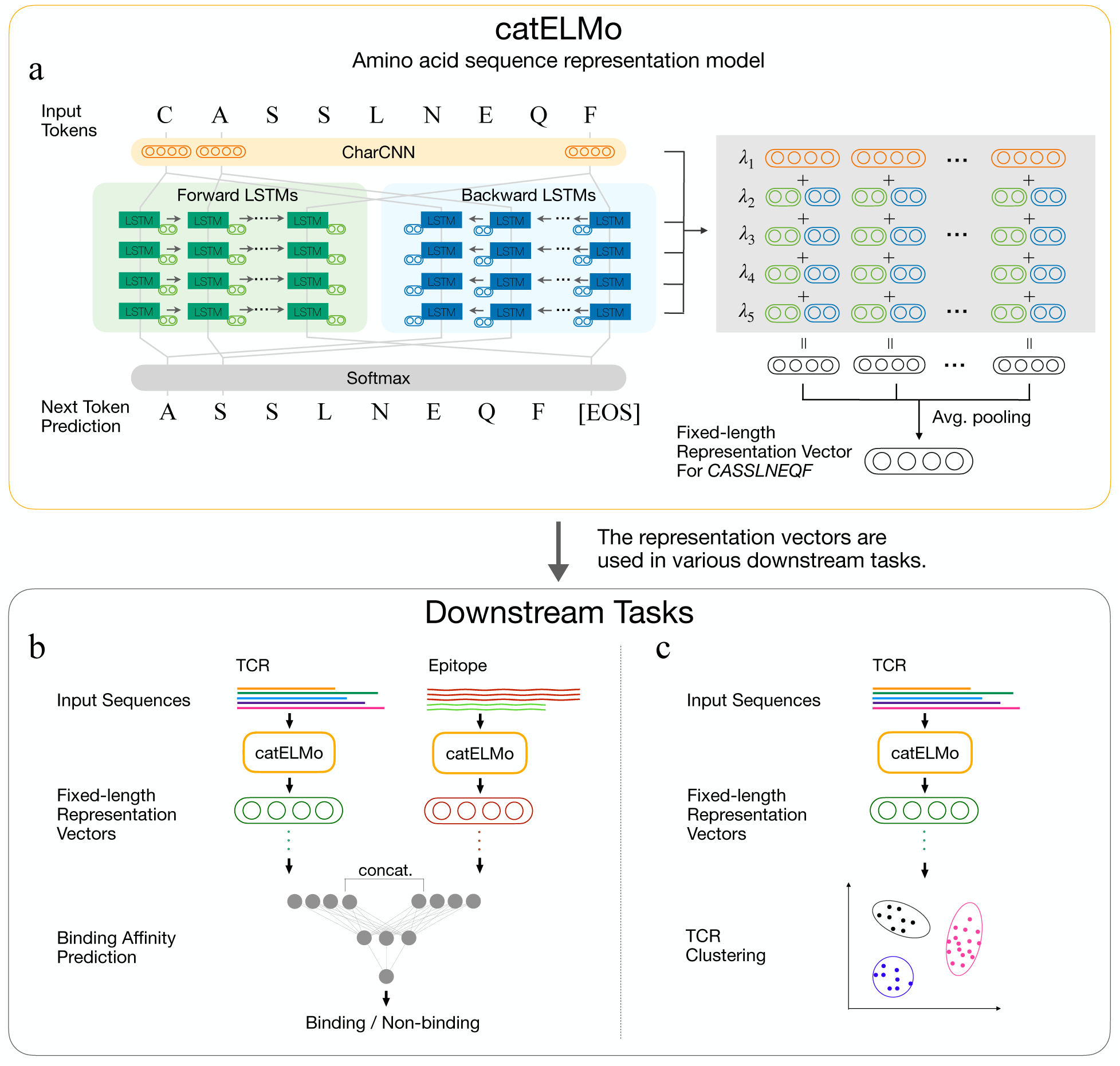
**Methods overview. a**) `catELMo` is an ELMo-based bi-directional amino acid sequence representation model trained on TCR sequences. It takes a sequence of amino acid strings as input and predicts the right (or left) next amino acid tokens. `catELMo` consists of a charCNN layer and four bidirectional LSTM layers followed by a softmax activation. For a given TCR sequence of length L, each layer returns L vectors of length 1,024. The size of an embedded TCR sequence, therefore, is [5, L, 1024]. Global average pooling with respect to the length of TCR, L, is applied to get a representation vector with a size of 1, 024. **b**) TCR-epitope binding affinity prediction task. An embedding method (e.g., `catELMo`) is applied to both TCR and epitope sequences. The embedding vectors are then fed into a neural network of three linear layers for training a binding affinity prediction model. The model predicts whether a given TCR and epitope sequence bind or not. **c**) Epitope-specific TCR sequences clustering. The hierarchical clustering algorithm is applied to TCR embedding vectors to group TCR sequences based on their epitope-specificity.

**Table 1.** Data Summary. The number of unique epitopes, TCRs, and TCR-epitope pairs used for `catELMo` and downstream tasks analysis.

| Usage | Source | Unique Epitopes | Unique TCRs | Unique TCR-epitope Pairs | Amino Acid Tokens |
| --- | --- | --- | --- | --- | --- |
| catELMo Training | ImmunoSEQ [23] | ✗ | 4,173,895 | ✗ | 52,546,029 |
| Binding Affinity Prediction | VDJdb* [33] | 187 | 3,915 | 4,047 | ✗ |
|  | McPAS [30] | 301 | 9,822 | 10,156 | ✗ |
|  | IEDB [34] | 1189 | 136,492 | 145,678 | ✗ |
|  | Total (after removing duplicates) | 982 | 140,675 | 150,008 | ✗ |
| Epitope-specific TCR Clustering | McPAS [30] Human, Mice | 8 | 5,607 | 5,607 | ✗ |
|  | McPAS [30] Human | 8 | 5,528 | 5,528 | ✗ |
|  | McPAS [30] Mice | 8 | 1,322 | 1,322 | ✗ |

\*VDJdb pairs with a score larger than 0 were used. ✗ represents not applicable. For binding affinity prediction task, we additionally generated 150,008 unique non-binding TCR-epitope sequences as negative samples.

We briefly summarize the two downstream tasks here and refer further details to Section 4.4. The first downstream task is TCR-epitope binding affinity prediction (Fig. 1b). All embedding models compared were to embed input sequences into the identical prediction model. Each prediction model was trained on 300,016 TCR-epitope binding and non-binding pairs (1:1 ratio), embedded by each embedding model. We used a neural network with three linear layers for the prediction model, which takes a pair of TCR and epitope as input and returns a binding affinity (0–1) of the pair. The prediction performance was evaluated on testing sets each defined by two types of splitting methods [10], called TCR and epitope splits. The testing set of TCR split has no TCRs overlapped with training and validation sets, allowing us to measure out-of-sample TCR performance. Similarly, the testing set of epitope split has no epitopes overlapped with training and validation sets, allowing us to measure out-of-sample epitope performance. For a fair comparison, a consistent embedding method was applied to both TCR and epitope sequences within a single prediction model. The second task is epitope-specific TCR clustering that aims at grouping TCRs that bind to the same epitope (Fig. 1c). We tested with TCR sequences of human and mouse species sampled from McPAS [30] database.

We applied the hierarchical clustering [31] and reported normalized mutual information (NMI) [32] and cluster purity to qualify the goodness of the clustering partition of TCR sequences.

### 2.1 catELMo outperforms the existing embedding methods at discriminating binding and non-binding TCR-epitope pairs

We investigated the downstream performance of TCR-epitope binding affinity prediction models trained using `catELMo` embeddings. In order to compare performance across different embedding methods, we used the identical downstream model architecture for each method. The competing embedding methods compared are BLOSUM62 [16], Yang et al. [25], ProBert [27], SeqVec [26] and TCRBert [29]. We observed that the prediction model using `catELMo` embeddings significantly outperformed those using existing amino acid embedding methods in both TCR (Figure 2a, b) and epitope (Figure 2d, e) split. In TCR split, where no TCRs in the testing set exist in the training and validation set, `catELMo`’s prediction performance was significantly greater than the second best method (p-value < 6.28 × 10^−23^, Table 2). It achieved AUC 96.04% which was 14% points higher than that of the second-highest performing method, while the rest of the methods performed worse than or similar to BLOSUM62. In epitope split, where no epitopes in the testing set exist in the training and validation set, the prediction model using `catELMo` also outperformed others with even larger performance gaps. `catELMo` significantly boosted 17% points of AUCs than the second-highest performing method (p-value < 1.18 × 10^−7^, Table 3). Similar performance gains from `catELMo` were also observed in other metrics such as Precision, Recall, and F1 scores (Supplementary Fig. 1).

**Figure 2.**
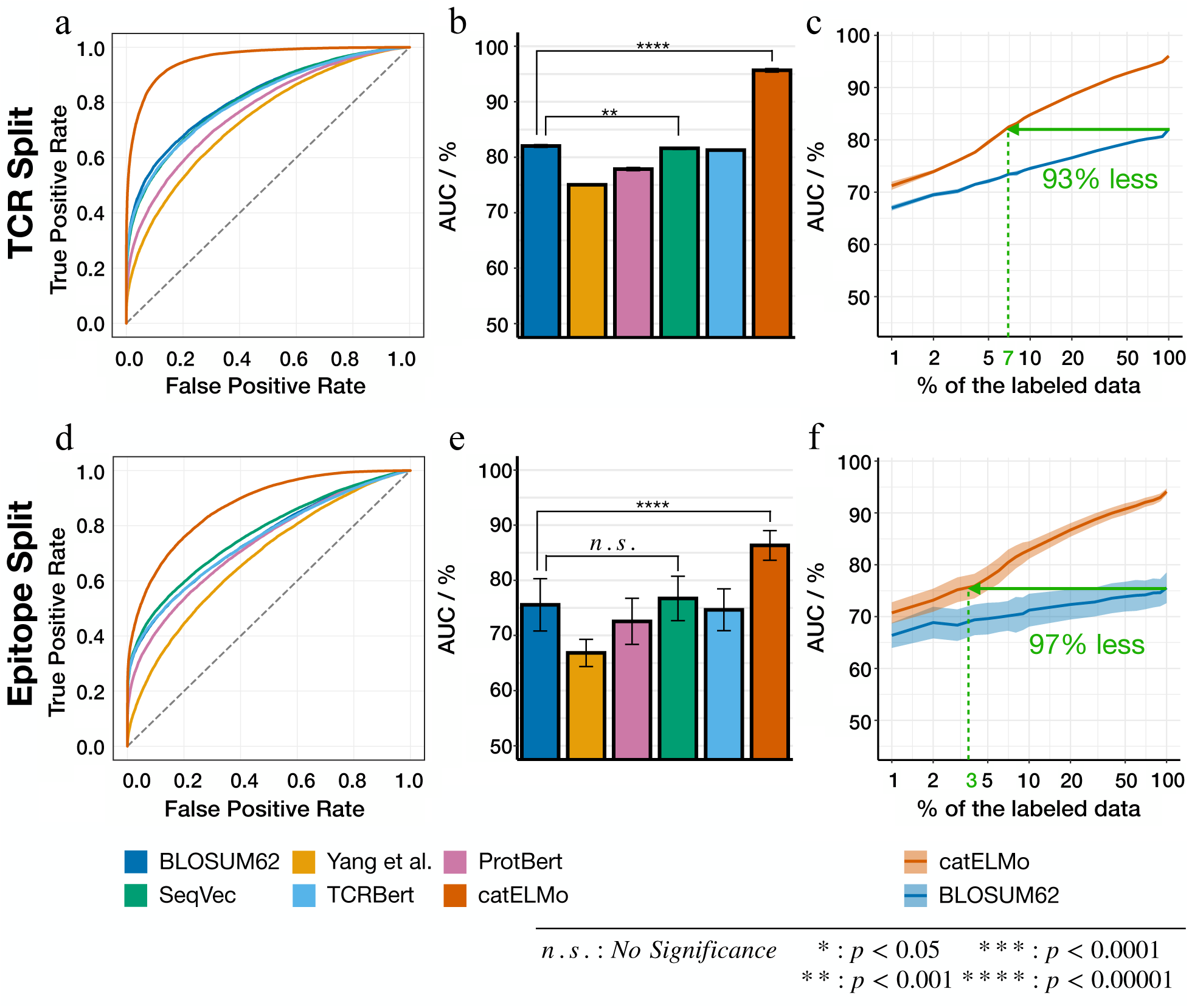
Comparison of the amino acid embedding methods for TCR-epitope binding affinity prediction task. We obtained TCR and epitope embeddings and used them as input features of binding affinity prediction model. A binding affinity prediction model is trained on each embedding method’s embedding dataset. The prediction performance comparison on **a), b), c)** TCR split and **d), e), f)** epitope split. **a), d)** Receiver Operating Characteristic (ROC) curve and **b), e)** AUC of the prediction model trained on different embedding methods. Error bars represent standard deviations over 10 trials. **c), f)** AUCs of the prediction model trained on different portions of downstream datasets. Error bands represent 95% confidence intervals over 10 trials.

**Table 2.** TCR-epitope binding affinity prediction performance of TCR split. Average and standard deviation of 10 trials are reported. P-values are from two-sample t-tests between `catELMo` and the second best method (<u>underlined</u>).

|  | AUC (%) | Precision (%) | Recall (%) | F1 (%) |
| --- | --- | --- | --- | --- |
| BLOSUM62 | <u>82.03 ± 0.25</u> | 67.16 ± 1.01 | <u>82.04 ± 1.01</u> | 70.57 ± 0.73 |
| Yang et al. | 75.03 ± 0.20 | 62.54 ± 0.78 | 79.71 ± 1.45 | 65.22 ± 0.69 |
| ProtBert | 77.86 ± 0.29 | 70.01 ± 1.47 | 69.90 ± 2.65 | 69.85 ± 0.41 |
| SeqVec | 81.61 ± 0.21 | 69.30 ± 1.33 | 79.02 ± 2.02 | 71.75 ± 0.66 |
| TCRBert | 80.79 ± 0.17 | <u>74.19 ± 1.17</u> | 70.48 ± 1.60 | <u>72.89 ± 0.23</u> |
| catELMo (ours) | **96.04 ± 0.12** | **86.88 ± 0.92** | **91.83 ± 0.98** | **88.94 ± 0.21** |
| p-value | $6.28 \times 10^{-23}$ | $1.94 \times 10^{-15}$ | $1.82 \times 10^{-14}$ | $1.29 \times 10^{-29}$ |

**Table 3.** TCR-epitope binding affinity prediction performance of epitope split. Average and standard deviation of 10 trials are reported. P-values are from two-sample t-tests between `catELMo` and the second best method (<u>underlined</u>).

|  | AUC (%) | Precision (%) | Recall (%) | F1 (%) |
| --- | --- | --- | --- | --- |
| BLOSUM62 | 75.54 ± 4.74 | 64.87 ± 2.17 | 73.12 ± 5.72 | 66.65 ± 2.94 |
| Yang et al. | 66.83 ± 2.46 | 58.68 ± 1.19 | 73.00 ± 3.65 | 60.16 ± 1.52 |
| ProtBert | 72.55 ± 4.18 | 67.66 ± 3.12 | 62.21 ± 6.93 | 66.04 ± 3.23 |
| SeqVec | <u>76.71 ± 4.02</u> | 66.35 ± 2.31 | <u>73.49 ± 6.26</u> | <u>67.92 ± 2.74</u> |
| TCRBert | 74.43 ± 3.98 | <u>73.34 ± 2.07</u> | 57.02 ± 7.70 | 67.64 ± 3.45 |
| catELMo (ours) | **94.10 ± 0.90** | **84.64 ± 1.39** | **88.85 ± 2.04** | **86.33 ± 1.04** |
| p-value | $1.18 \times 10^{-7}$ | $1.89 \times 10^{-10}$ | $1.47 \times 10^{-5}$ | $2.71 \times 10^{-10}$ |

We also visually observed that `catELMo` aided the model to better discriminate binding and non-binding TCRs for the five most frequent epitopes that appeared in our collected TCR-epitope pairs (Fig. 3). These five epitopes account for a substantial portion of our dataset, comprising 14.73% (44,292 pairs) of the total TCR-epitope pairs collected. For visualization, we performed t-SNE [35] on the top fifty principal components of the last latent vectors of each prediction model. Each point represents a pair of TCR-epitope, colored by epitope (lighter shade for positive binding and darker shade for negative binding). Different degrees of overlapping between positive pairs and negative ones in regard to the same epitope can be seen in the t-SNE plots. For example, most of the binding and non-binding data points from SeqVec embeddings are barely separated within each epitope group. On the other hand, the t-SNE plot of `catELMo` exhibits noticeable contrast between binding and non-binding pairs, indicating that `catELMo` aids the prediction model to distinguish labels. We also observed `catELMo` outperformed the other embedding methods in discriminating binding and non-binding TCRs for almost all individual epitopes. As shown in Supplementary Fig. 2, the prediction model using `catELMo` embeddings achieved the highest AUCs in 39 out of 40 epitopes, and the second highest AUC on an epitope (`GTSGSPIVNR`) with only 1.09% lower than the highest score. Addtionally, we observed that `catELMo` consistently outperformed other embedding methods in predicting the binding affinity between TCRs and epitopes from a diverse range of pathogens (Supplementary Table 1).

**Figure 3.**
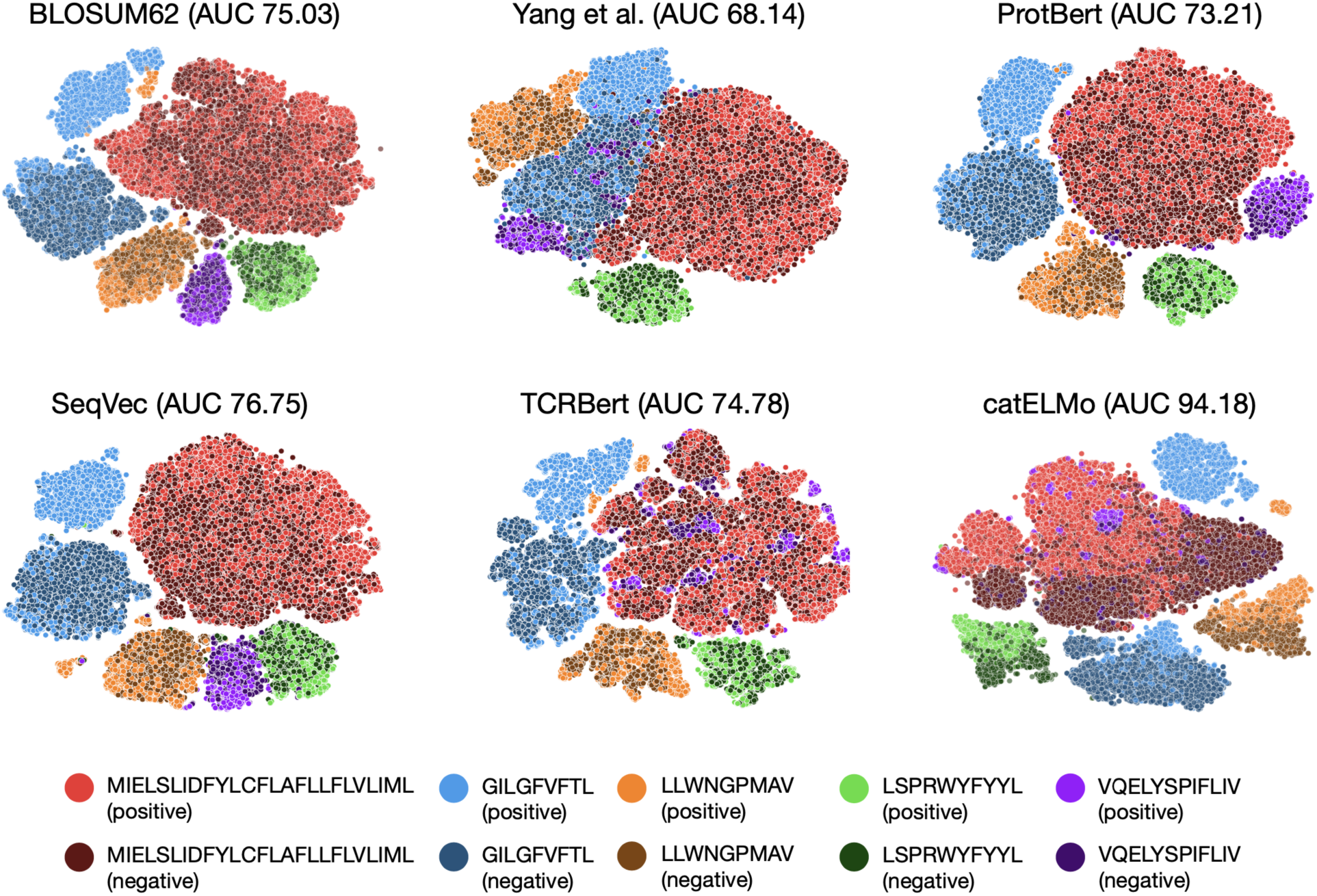
tSNE visualization for top five frequent epitopes. We visually compared the embedding models on TCR-epitope binding affinity prediction task. We conduct tSNE analysis on the top 50 principle components of the last hidden layer features of the TCR-epitope binding affinity prediction models for out-of-sample epitopes. The clearer the boundary between positive pairs (lighter shade) and negative pairs (darker shade) associated with the same epitope sequence, the better the model is at discriminating binding and non-binding pairs.

### 2.2 catELMo reduces a significant amount of annotation cost for achieving comparable prediction power

Language models trained on large corpus are known to improve downstream task performance with a smaller number of downstream training data [14, 19]. Similarly in TCR-epitope binding, we show that `catELMo` trained entirely on unlabeled TCR sequences facilitates its downstream prediction model to achieve the same performance with a significantly smaller amount of TCR-epitope training pairs (i.e., epitope-labeled TCR sequence). We trained a binding affinity prediction model for each k% of downstream data (i.e., `catELMo` embeddings of TCR-epitope pairs) where k = 1, 2, ···, 10, 20, 30, ···, 100. The widely used BLOSUM62 embedding matrix was used as a comparison baseline under the same ks as it performs better than or is comparable to the other embedding methods.

A positive log-linear relationship between the number of (downstream) training data and AUCs was observed for both TCR and epitope split (Fig. 2c, f). The steeper slope in `catELMo` suggests that prediction models utilizing `catELMo` embeddings exhibit higher performance gain per number of training pairs compared to the BLOSUM62-based models. In TCR split, we observed that `catELMo`’s binding affinity prediction models with just 7% of the training data significantly outperform ones that use a full size of BLOSUM62 embeddings (p-value = 0.0032, Fig. 2c). `catELMo` with just 3%, 4%, and 6% of the downstream training data achieved similar performances to when using a full size of Yang et al., ProtBert, and SeqVec embeddings, respectively. Similarly, in epitope split, we showed `catELMo`’s prediction models with just 3% of training data achieved equivalent performance as ones built on a full size of BLOSUM62 embeddings (p-value = 0.8531, Fig. 2f). Compare to the other embedding methods, `catELMo` with just 1%, 2%, and 5% of the downstream training data achieved similar or better performance than when using a full size of Yang et al., ProtBert, and SeqVec embeddings, separately. Similar performance gains from `catELMo` were also observed in other metrics such as Precision, Recall, and F1 scores (Supplementary Fig. 3). Achieving accurate prediction with a small amount of training data is important for TCR analysis as obtaining the binding affinity of TCR-epitope pairs is costly.

### 2.3 catELMo allows clustering of TCR sequences with high performance

Clustering TCRs of similar binding profiles is important in TCR repertoire analysis as it facilitates discoveries of TCR clonotypes that are condition-specific [36, 37]. In order to demonstrate that `catELMo` embeddings can be used for other TCR-related downstream tasks, we performed hierarchical clustering [31] using each method’s embedding (`catELMo`, BLOSUM62 [16], Yang et al. [25], ProBert [27], SeqVec [26] and TCRBert [29]) and evaluated the identified clusters against the ground-truth TCR groups labeled by their binding epitopes. We additionally compared our results with state-of-the-art TCR clustering methods, TCRdist [37] and GIANA [38], both of which were developed from BLOSUM62 matrix (see Section 4.4.2). Normalized mutual information (*NMI*) [32] and cluster purity are used to measure the clustering quality.

Significant disparities in TCR binding frequencies exist across different epitopes. To construct more balanced clusters, we targeted TCR sequences bound to the top eight frequent epitopes identified in the McPAS database. We find that the cluster model for both human and mouse species built on `catELMo` embeddings maintains either the best or second best NMI and purity scores compared with ones that are computed on other embeddings (Fig. 4a, d). To investigate whether this observation remains true on individual species, we conducted the same clustering analysis on human and mouse species, separately. We showcase comparison for the top eight epitopes in human (Fig. 4b, e) and mouse (Fig. 4c, f) species and observe a similar pattern that clustering results with `catELMo` achieve the highest or second-highest NMI and purity scores. Since the real-world scenarios likely involve more epitopes, we conducted additional clustering experiments using the top most abundant epitopes whose combined cognate TCRs make up at least 70% of TCRs across three databases (34 epitopes). Similar performance gains were observed (Supplementary Fig. 4). Altogether, `catELMo` embedding can assist TCR clustering with no supervision while achieving similar or better performance than other state-of-the-art methods in both human and mouse species.

**Figure 4.**
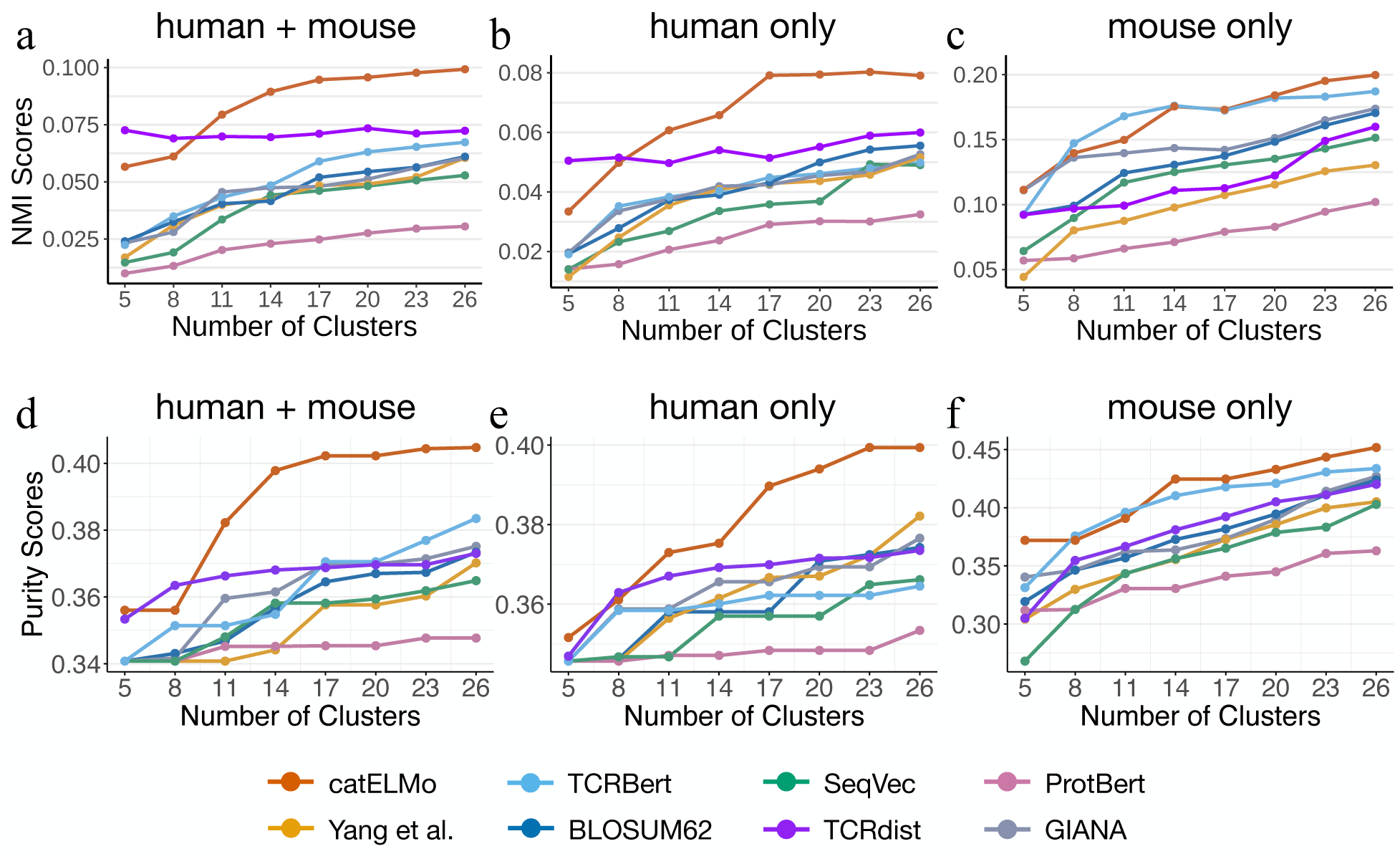
Comparison of the amino acid embedding methods for epitope-specific TCR clustering. Hierarchical clustering is applied on the McPAS [30] database. We cluster TCRs binding to the top eight epitopes of **a**) both human and mouse species, **b**) only human epitopes, and **c**) only mouse epitopes. Larger NMI scores indicate TCR sequences that bind to the same epitope are grouped together in the same cluster and TCR sequences that do not bind to the same epitope are separated apart in different clusters.

### 2.4 ELMo-based architecture is preferable to BERT-based architecture in TCR embedding models

We observed `catELMo` using ELMo-based architecture outperformed the model using embeddings of TCRBert which uses BERT (Table 4). The performance differences were approximately 15% AUCs in TCR split (p-value < 3.86 × 10^−30^) and 19% AUCs in epitope split (p-value < 3.29 × 10^−8^). Because TCRBert was trained on a smaller amount of TCR sequences (around 0.5 million sequences) than `catELMo`, we further compared `catELMo` with various sizes of BERT-like models trained on the same dataset as `catELMo`: BERT-Tiny-TCR, BERT-Base-TCR, and BERT-Large-TCR having a stack of 2, 12, and 30 Transformer layers respectively (see Section 4.6.2). Note that BERT-Base-TCR uses the same number of Transformer layers as TCRBert. Additionally, we compared different versions of `catELMo` by varying the number of BiLSTM layers (2, 4–default, and 8, see Section 4.6.1). As summarized in Table 4, TCR-epitope binding affinity prediction models trained on `catELMo` embeddings (AUC 96.04% and 94.70% on TCR and epitope split) consistently outperformed models trained on these Transformer-based embeddings (AUC 81.23–81.91% and 74.20–74.94% on TCR and epitope split). The performance gaps between `catELMo` and Transformer-based models (14% AUCs in TCR split and 19% AUCs in epitope split) were statistically significant (p-values < 6.72 × 10^−26^ and < 1.55 × 10^−7^ for TCR and epitope split respectively). We observed that TCR-epitope binding affinity prediction models trained on `catELMo`-based embeddings consistently outperformed the ones using Transformer-based embeddings (Table 4, 5). Even the worst- performed BiLSTM-based embedding model achieved higher AUCs than the best-performed Transformer-based embeddings at discriminating binding and non-binding TCR-epitope pairs in both TCR (p-value < 2.84 × 10^−28^) and epitope split (p-value < 5.86 × 10^−6^).

**Table 4.**
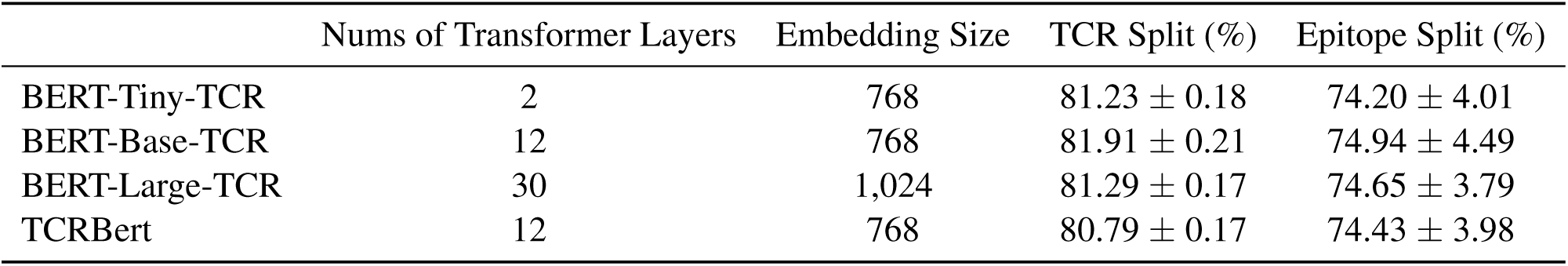
AUCs of TCR-epitope binding affinity prediction models built on BERT-based embedding models. Average and standard deviation of 10 trials are reported.

|  | Nums of Transformer Layers | Embedding Size | TCR Split (%) | Epitope Split (%) |
| --- | --- | --- | --- | --- |
| BERT-Tiny-TCR | 2 | 768 | $81.23 \pm 0.18$ | $74.20 \pm 4.01$ |
| BERT-Base-TCR | 12 | 768 | $81.91 \pm 0.21$ | $74.94 \pm 4.49$ |
| BERT-Large-TCR | 30 | 1,024 | $81.29 \pm 0.17$ | $74.65 \pm 3.79$ |
| TCRBert | 12 | 768 | $80.79 \pm 0.17$ | $74.43 \pm 3.98$ |

**Table 5.** AUCs of TCR-epitope binding affinity prediction models trained on different sizes of catELMo embeddings. Average and standard deviation of 10 trials are reported.

|  | Nums of BiLSTM Layers | Embedding Size | TCR Split (%) | Epitope Split (%) |
| --- | --- | --- | --- | --- |
| catELMo-Shallow | 2 | 1,024 | $95.67 \pm 0.32$ | $86.32 \pm 2.68$ |
| catELMo | 4 | 1,024 | **$96.04 \pm 0.12$** | **$94.10 \pm 0.90$** |
| catELMo-Deep | 8 | 1,024 | $93.94 \pm 0.19$ | $91.57 \pm 1.59$ |

### 2.5 Within-domain transfer learning is preferable to cross-domain transfer learning in TCR analysis

We showed that `catELMo`, trained on TCR sequences, significantly outperformed amino acid embedding methods trained on generic protein sequences. `catELMo`-Shallow and SeqVec shared the same architecture consisting of character-level convolutional layers and a stack of two bi-directional LSTM layers but were trained on different types of training data. `catELMo`-Shallow was trained on TCR sequences (about 4 million) while SeqVec was trained on generic protein sequences (about 33 million). Although `catELMo`-Shallow was trained on a relatively smaller amount of sequences compared to SeqVec, the binding affinity prediction model built on `catELMo`-Shallow embeddings (AUC 95.67% in TCR split and 86.32% in epitope split) significantly outperformed the one built on SeqVec embeddings (AUC 81.61% in TCR split and 76.71% in epitope split) by 14.06% and 9.61% on TCR and epitope split respectively. This suggests that knowledge transfer within the same domain is preferred whenever possible in TCR analysis.

## 3 Discussion

`catELMo` is an effective embedding model that brings substantial performance improvement in TCR-related downstream tasks. Our study emphasizes the importance of choosing the right embedding models. The embedding of amino acids into numeric vectors is the very first and crucial step that enables the training of a deep neural network. It has been previously demonstrated that a well-designed embedding can lead to significantly improved results on downstream analysis [25, 26, 27]. The reported performance of `catELMo` embedding on TCR-epitope binding affinity prediction and TCR clustering tasks indicates that `catELMo` is able to learn patterns of amino acid sequences more effectively than state-of-the-art embedding methods. While all other methods compared (except BLOSUM62) leverage a large number of unlabeled amino acid sequences, only our prediction model using `catELMo` significantly outperforms widely used BLOSUM62 and other models such as netTCR [9, 17] and ATM-TCR [10] trained on paired (TCR-epitope) samples only (Table 6). Our work suggests the need for developing sophisticated strategies to train amino acid embedding models that can enhance the performance of TCR-related downstream tasks even while requiring less amount of data and simpler prediction model structures.

**Table 6.**
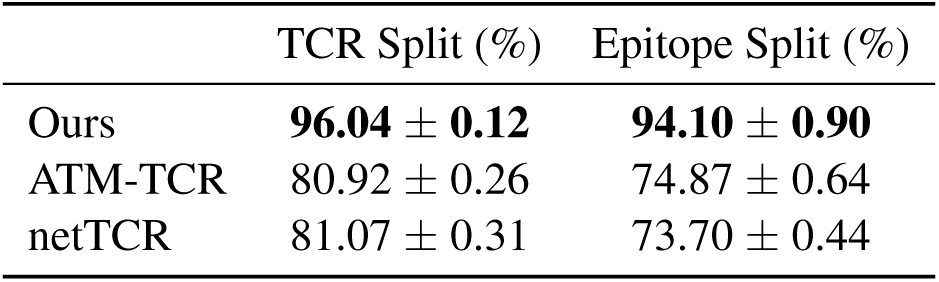
AUCs of TCR-epitope binding affinity prediction models comparison with state-of-the-art prediction models. All models are trained on the same dataset. Average and standard deviation of 10 trials are reported.

|  | TCR Split (%) | Epitope Split (%) |
| --- | --- | --- |
| Ours | **96.04 ± 0.12** | **94.10 ± 0.90** |
| ATM-TCR | 80.92 ± 0.26 | 74.87 ± 0.64 |
| netTCR | 81.07 ± 0.31 | 73.70 ± 0.44 |

Two important observations made from our experiments are 1) the type of data used for training amino acid embedding models is far more important than the amount of data and 2) ELMo-based embedding models consistently perform much better than BERT-based embedding models. While previously developed amino acid embedding models such as SeqVec and ProtBert were trained on 184- and 1,690-times more amino acid tokens compared to the training data used for `catELMo`, the prediction models using SeqVec and ProtBert performed poorly compared to the model using `catELMo` (see Sections 2.1 and 2.3). SeqVec and ProtBert were trained based on generic protein sequences, whereas `catELMo` was trained on a collection of TCR sequences from pooled TCR repertoires across many samples, indicating that the use of TCR data to train embedding models is more critical than much larger amount of generic protein sequences.

In the field of natural language processing, Transformer-based models have been bolstered as the superior embedding model [14, 15, 19]. However, for TCR-related downstream tasks, `catELMo` using biLSTM layer-based design outperforms BERT using Transformer layers (see Section 2.4). While it is difficult to pinpoint the reasons, the bi-directional architecture to predict the next token based on its previous tokens in ELMo may mimic the interaction process of TCR and epitope sequences either from left to right or from right to left. In contrast, BERT uses Transformer encoder layers that attend tokens both on the left and right to predict a masked token, refer to as masked language modeling. As the Transformer layer can be along with the next token prediction objectives, it remains as a future work to investigate Transformer with causal language models, such as GPT-3 [14], for amino acid embedding. Additionally, the clear differences of TCR sequences compared to natural languages are 1) the compact vocabulary size (20 standard amino acids vs. over 170k English words) and 2) the length of peptides in TCRs being smaller than the number of words in sentences or paragraphs in natural languages. These differences may allow `catELMo` to learn sequential dependence without losing long-term memory from the left end.

Often in classification problems in life sciences, the difference in the number of available positive and negative data can be very large and TCR-epitope binding affinity prediction problem is no exception. In fact, the number of experimentally generated non-binding pairs are practically non-existent and obtaining experimental negative data is costly [17]. This requires researchers to come up with a strategy to generate negative samples and it can be non-trivial. A common practice is to sample new TCRs from repertoires and pair them with existing epitopes [9, 39], a strategy we also used. Another approach is to randomly shuffle TCR-epitope pairs within positive binding dataset, resulting in TCRs and epitopes that are not known to bind paired together [10, 11]. Given the vast diversity of human TCR clonotypes, which can exceed 10^15^ [40], the chance of randomly selecting a TCR that specifically recognizes a target epitope is relatively small. The prediction model consistently outperformed the other embedding methods by large margins in both TCR and epitope splits as shown in Supplementary Fig. 5. The model using `catELMo` achieves 24% and 36% higher AUCs over the second best embedding method for TCR (p-value < 1.04 × 10^−18^, Supplementary Table 2) and epitope (p-value < 6.26 × 10^−14^, Supplementary Table 3) split, respectively. Moreover, we observe that using `catELMo` embeddings, prediction models that are trained with only 2% downstream samples still statistically outperform ones that are built on a full size of BLOSUM62 embeddings in TCR split (p-value = 0.0005). Similarly, with only 1% training samples, `catELMo` reaches comparable results as BLOSUM62 with a full size of downstream samples in epitope split (p-value = 0.1438). In other words, `catELMo` dramatically reduces about 98% annotation cost (Supplementary Table 4). To mitigate potential batch effects, we generated new negative pairs using different seeds and observed consistent prediction performance across these variations. Our experiment results confirm that the embeddings from `catELMo` maintain high performance regardless of the methodology used to generate negative samples.

Parameter fine-tuning in neural networks is a training scheme where initial weights of the network are set to the weights of a pre-trained network. Fine-tuning has been shown to bring performance gain to the model over using random initial weights [41]. We investigated the possibility of performance boost of our prediction model using fine-tuned `catELMo`. Since SeqVec shares the same architecture with `catELMo`-Shallow and is trained on generic protein sequences, we used the weights of SeqVec as initial weights when fine-turning `catELMo`-Shallow. We compared the performance of binding affinity prediction models using the fine-tuned `catELMo`-Shallow and vanilla `catELMo`-Shallow (trained from scratch with random initial weights from a standard normal distribution). We observed that the performance when using fine-tuned `catELMo`-Shallow embeddings was significantly improved by approximately 2% AUCs in TCR split (p-value < 4.17 × 10^−9^) and 9% AUCs in epitope split (p-value < 5.46 × 10^−7^).

While epitope embeddings are a part of our prediction models, their impact on overall performance appears to be less significant compared to that of TCR embeddings. To understand the contribution of epitope embeddings, we performed additional experiments. First, we kept epitope embeddings unchanged using the widely-used BLOSUM62-based while varying different embedding methods exclusively for TCRs. The results (Supplementary Table 5) closely aligns with our previous findings (Tables 2 and 3), suggesting that the choice of epitope embedding method may not strongly affect the final predictive performance.

Furthermore, we investigated alternative embedding approaches for epitope sequences. Specifically, we replaced epitope embeddings with randomly initialized matrices containing trainable parameters, while employing `catELMo` for TCR embeddings. This setting yielded predictive performance comparable to the scenario where both TCR and epitope embeddings were `catELMo`-based (Supplementary Table 6).

Similarly, using BLOSUM62 for TCR embeddings and `catELMo` for epitope embeddings resulted in performance similar to when both embeddings were based on BLOSUM62. These consistent findings support the proposition that the influence of epitope embeddings may not be as significant as that of TCR embeddings (Supplementary Table 7).

We believe these observations may be attributed to the substantial data scale discrepancy between TCRs (more than 290k) and epitopes (less than 1k). Moreover, TCRs tend to exhibit high similarity, whereas epitopes display greater distinctiveness from one another. These features of TCRs require robust embeddings to facilitate effective separation and improve downstream performance, while epitope embeddings primarily serve as categorical encodings.

While TCR*β* CDR3 is known to be the primary determinant for TCR-epitope binding specificity, other regions such as CDR1 and CDR2 on TCR*β* V gene along with TCRα chain are also known to contribute to specificity in antigen recognition [42, 43].There are models that can take advantage of these additional features when making predictions [17, 44]. However, our study focuses on modeling CDR3 of TCR*β* chains because of the limited availability of sample data from other regions. Future work may explore strategies to incorporate these regions while mitigating the challenges of working with limited samples.

## 4 Methods

We first present data used for training the amino acid embedding models and the downstream tasks. We then review existing amino acid embedding methods and their usage on TCR-related tasks. We introduce our approach, `catELMo`, a bi-directional amino acid embedding method that computes contextual representation vectors of amino acids of a TCR (or epitope) sequence. We describe in detail how to apply `catELMo` to two different TCR-related downstream tasks. Lastly, we provide details on the experimental design, including the methods and parameters used in comparison and ablation studies.

### 4.1 Data

#### TCRs for training catELMo

We collected 5,893,249 TCR sequences from repertoires of seven projects in the ImmunoSEQ database: HIV [45], SARS-CoV2 [23], Epstein Barr Virus [46], Human Cytomegalovirus [47], Influenza A [48], Mycobacterium Tuberculosis [49], and Cancer Neoantigens [50]. CDR3 sequences of TCR*β* chains were used to train the amino acid embedding models as those are the major segment interacting with epitopes [2] and exist in large numbers. We excluded duplicated copies and sequences containing wildcards such as ‘*’ or ‘X’. Altogether, we obtained 4,173,895 TCR sequences (52,546,029 amino acid tokens) of which 85% were used for training and 15% were used for testing.

#### TCR-epitope pairs for binding affinity prediction

We collected TCR-epitope pairs known to bind each other from three publicly available databases: IEDB [34], VDJdb [33], and McPAS [30]. Unlike the (unlabeled) TCR dataset for `catELMo` training, each TCR is annotated with an epitope known to bind each other, which we referred to as a TCR-epitope pair. We only used pairs with human MHC class I epitopes and CDR3 sequences of the TCR*β* chain. We further filtered out sequences containing wildcards such as ‘*’ or ‘X’. For VDJdb, we excluded pairs with a confidence score of 0 as it means a critical aspect of sequencing or specificity validation is missing. We removed duplicated copies and merged datasets collected from the three databases. For instance, 29.85% of pairs from VDJdb overlapped with IEDB, and 55.41% of pairs from McPAS overlapped with IEDB. Altogether, we obtained 150,008 unique TCR-epitope pairs known to bind to each other having 140,675 unique TCRs and 982 unique epitopes. We then generated the same number of non-binding TCR-epitope pairs as negative samples by randomly pairing each epitope of the positive pairs with a TCR sampled from the healthy TCR repertoires of ImmunoSEQ. We note that it includes no identical TCR sequences with the TCRs used for training the embedding models. Altogether, we obtained 300,016 TCR-epitope pairs where 150,008 pairs are positive and 150,008 pairs are negative. The average length of TCRs and epitope sequences are 14.78 and 11.05, respectively. Our data collection and preprocessing procedures closely followed those outlined in our previous work [10].

#### TCRs for antigen-specific TCR clustering

We collected 9,822 unique TCR sequences of humans and mice hosts from McPAS [30]. Each TCR is annotated with an epitope known to bind, which is used as a ground-truth label for TCR clustering. We excluded TCR sequences that bind to neoantigen pathogens or multiple epitopes and only used CDR3 sequences of TCR*β* chain. We composed three subsets for different experimental purposes. The first dataset contains both human and mice TCRs. We used TCRs associated with the top eight frequent epitopes, resulting in 5,607 unique TCRs. The second dataset consists of only human TCRs, and the third dataset consists of only mouse TCRs. In a similar manner, we selected TCRs that bind to the top eight frequent epitopes. As a result, we obtained 5,528 unique TCR sequences for the second dataset and 1,322 unique TCR sequences for the third dataset.

### 4.2 Amino acid embedding methods

In this section, we review amino acid embedding methods previously proposed. There are two categories of the existing approaches: static and context-aware embedding methods. Static embedding method represents an amino acid as a static representation vector remaining the same regardless of its context. Context-aware embedding method, however, represents an amino acid differently in accordance with its context. Context-aware embedding is also called dynamic embedding in contrast to static embedding. We explain the key ideas of various embedding methods, and introduce their usage in previous works.

#### 4.2.1 Static embeddings

##### BLOSUM

BLOSUM [16] is a scoring matrix where each element represents how likely an amino acid residue is to be substituted by another over evolutionary time. It has been commonly used to measure alignment scores between two protein sequences. There are various BLOSUM matrices such as BLOSUM45, BLOSUM62, and BLOSUM80 where a matrix with a higher number is used for the alignment of less divergent sequences. BLOSUM have also served as the de facto standard embedding method for various TCR analyses. For example, BLOSUM62 was used to embed TCR and epitope sequences for training deep neural network models predicting their binding affinity [7, 9, 17]. BLOSUM62 was also used to embed TCR sequences for antigen-specific TCR clustering and TCR repertoire clustering. GIANA [38] clustered TCRs based on the Euclidean distance between TCR embeddings. TCRdist [37] used BLOSUM62 matrix to compute the dissimilarity matrix between TCR sequences for clustering.

##### Word2vec and Doc2vec

Word2vec [20] and Doc2vec [21] are a family of embedding models to learn a single linear mapping of words, which takes a one-hot word indicator vector as input and returns a real-valued word representation vector as output. There are two types of Word2vec architectures: continuous bag-of-words (CBOW) and skip-gram. CBOW predicts a word from its surrounding words in a sentence. It embeds each input word via a linear map, sums all input words’ representations, and applies a softmax layer to predict an output word. Once training is completed, the linear mapping is used to obtain a representation vector of a word. On the contrary, skip-gram predicts the surrounding words given a word while it also uses a linear mapping to obtain a representation vector. Doc2vec is a model further generalized from Word2vec, which introduces a paragraph vector representing paragraph identity as an additional input. Doc2vec also has two types of architectures: distributed memory (DM) and distributed bag-of-words (DBOW). DM predicts a word from its surrounding words and the paragraph vector, while DBOW uses the paragraph vector to predict randomly sampled context words. In a similar way, linear mapping is used to obtain a continuous representation vector of a word.

Several studies adapted Word2vec and Doc2vec to embed amino acid sequences [24, 51, 25]. ProtVec [24] is the first Word2vec representation model trained on a large number of amino acid sequences. Its embeddings were used for several downstream tasks such as protein family classification, disordered protein visualization, and classification. Kimothi et al. [51] adapted Doc2vec to embed amino acid sequences for protein sequence classification and retrieval. Yang et al. [25] trained Doc2vec models on 524,529 protein sequences of UniProt [22] database. They considered a k-mer amino acids as a word, and a protein sequence as a paragraph. They trained DM models to predict a word from w surrounding words and a paragraph with various sizes of k and w.

#### 4.2.2 Context-aware embeddings

##### ELMo

ELMo [18] is a deep context-aware word embedding model trained on a large corpus. It learns each token’s (e.g., a word) contextual representation in forward and backward directions using a stack of two bi-directional LSTM layers. Each word of a text string is first mapped into a numerical representation vector via the character-level convolutional layers. The forward (left-to-right) pass learns a token’s contextual representation depending on itself and the previous context in which it is used. The backward (left-to-right) pass learns a token’s representation depending on itself and its subsequent context.

ELMo is less commonly implemented for amino acid embedding than Transformer-based deep neural networks. One example is SeqVec [26]. It is an amino acid embedding model using ELMo’s architecture. It feeds each amino acid as a training token of size 1, and learns its contextual representation both forward and backward within a protein sequence. The data was collected from UniRef50 [52], which consists of 9 billion amino acid tokens and 33 million protein sequences. SeqVec was applied to several protein-related downstream tasks such as secondary structure and long intrinsic disorder prediction, and subcellular localization.

##### BERT

BERT [19] is a large language model leveraging Transformer [53] layers to learn context-aware word embeddings jointly conditioned on both directions. BERT is learned for two objectives. One is the masked language model to learn contextual relationships between words in a sentence. It aims to predict the original value of masked words. The other is the next sentence prediction which aims to learn the dependency between consecutive sentences. It feeds a pair of sentences as input and predicts whether the first sentence in the pair is contextually followed by the second sentence.

BERT’s architecture has been used in several amino acid embedding methods [27, 54, 29]. They treated an amino acid residue as a word and a protein sequence as a sentence. ProtBert [27] was trained on 216 million protein sequences (88 billion amino acid tokens) of UniRef100 [52]. It was applied for several protein sequence applications such as secondary structure prediction and sub-cellular localization. ProteinBert [54] combined language modeling and gene ontology annotation prediction together during training. It was applied to protein secondary structure, remote homology, fluorescence and stability prediction. TCRBert [29] was trained on 47,040 TCR*β* and 4,607 TCRα sequences of PIRD [55] dataset and evaluated on TCR-antigen binding prediction and TCR engineering tasks.

### 4.3 Our approach: catELMo

We propose `catELMo`, a bi-directional amino acid embedding model designed for TCR analysis. `catELMo` adapts ELMo’s architecture to learn context-aware representations of amino acids. It is trained on TCR sequences, which is different from the existing amino acid embedding models such as SeqVec trained on generic protein sequences. As illustrated in Fig. 1a, `catELMo` is composed of a character CNN (CharCNN) [56] layer converting each one-hot encoded amino acid token to a continuous representation vector, a stack of four bi-directional LSTM [57] layers learning contextual relationship between amino acid residues, and a softmax layer predicting the next (or previous) amino acid residue.

Given a sequence of N amino acid tokens, (*t*_1_, *t*_2_, …, *t_N_*), CharCNN maps each one-hot encoded amino acid token *t_k_* to a latent vector c*_k_* through seven convolutional layers with kernel sizes ranging from 1 to 7, and the numbers of filters of 32, 32, 64, 128, 256, 512, and 512, each of which is followed by a max-pooling layer, resulting in a 1,024-dimensional vector. The output of the CharCNN, (*c*_1_, *c*_2_, …, *c_N_*), is then fed into a stack of four bidirectional LSTM layers consisting of forward and backward passes. For the forward pass, the sequence of the CharCNN output is fed into the first forward LSTM layer followed by the second forward LSTM layer, and so on. Each LSTM cell in every forward layer has 4,096 hidden states and returns a 512-dimensional representation vector. Each output vector of the last LSTM layer is then fed into a softmax layer to predict the right next amino acid token. Residual connection is applied between the first and second layers and between the third and fourth layers to prevent gradient vanishing. Similarly, the sequence of the CharCNN output is fed into the backward pass, in which each cell returns a 512-dimensional representation vector. Unlike the forward layer, each output vector of the backward layer followed by a softmax layer aims to predict the left next amino acid token.

Through the forward and backward passes, `catELMo` models the joint probability of a sequence of amino acid tokens. The forward pass aims to predict the next right amino acid token given its left previous tokens, which is P (*t_k_*|*t*_1_, *t*_2_, …, *t_k—_*_1_; *θ_c_*, *θ_fw_*, *θ_s_*) for each k-th cell where *θ_c_* indicates parameters of CharCNN, *θ_fw_* indicates parameters of the forward layers, and *θ_s_* indicates parameters of the softmax layer. The joint probability of all amino acid tokens for the forward pass is defined as:

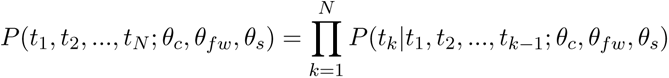

The backward pass aims to predict the next left amino acid token given its right previous tokens. Similarly, the joint probability of all amino acid tokens for the backward pass is defined as:

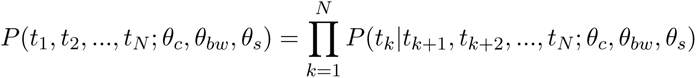

where *θ_bw_* indicates parameters of the backward layers. During `catELMo` training, the combined log-likelihood of the forward and backward passes is jointly optimized, which is defined as:

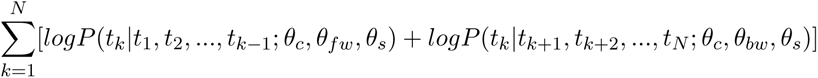

Note that the forward and backward layers have their own weights (*θ_fw_* and *θ_bw_*). This helps to avoid information leakage that a token used to predict its right tokens in forward layers is undesirably used again to predict its own status in backward layers.

For each amino acid residue, `catELMo` computes five representation vectors of length 1,024: one from CharCNN and four from BiLSTM layers. Those vectors are averaged over and yield an amino acid representation vector of length 1,024. A sequence of amino acids is then represented by an element-wise average of all amino acids’ representation vectors, resulting in a representation vector of length 1,024. For example, `catELMo` computes a representation for each amino acid in a TCR sequence CASSP T SGGQET QY F as a vector of length 1,024. The sequence is then represented by averaging over 15 amino acid representation vectors, which is a vector with a length of 1,024. `catELMo` is trained up to 10 epochs with a batch size of 128 on two NVIDIA RTX 2080 GPUs. We follow the default experimental settings of ELMo unless otherwise specified.

### 4.4 Downstream tasks

We evaluate the amino acid embedding models’ generalization performance on two downstream tasks: TCR-epitope binding affinity prediction and epitope-specific TCR clustering.

#### 4.4.1 TCR-epitope binding affinity prediction

Computational approaches that predict TCR-epitope binding affinity benefit rapid TCR screening for a target antigen and improve personalized immunotherapy. Recent computational studies [7, 9, 10, 17] formulated it as a binary classification problem that predicts a binding affinity score (0–1) given a pair of TCR and epitope sequences.

We evaluate `catELMo` based on the prediction performance of a binding affinity prediction model trained on its embedding, and compare it with the state-of-the-art amino acid embeddings (further demonstrated in Section 4.5). We first obtain different types of TCR and epitope embeddings from `catELMo` and the comparison methods. To measure the generalized prediction performance of binding affinity prediction models, we split each method’s dataset into training (64%), validation (16%), and testing (20%) sets. We use two splitting strategies established in the previous work [10]: TCR split and epitope split. TCR split was designed to measure the models’ prediction performance on out-of-sample TCRs where no TCRs in the testing set exist in the training and validation set. Epitope split was designed to measure the models’ prediction performance on out-of-sample epitopes where no epitopes in the testing set exist in the training and validation set.

The downstream model architecture is the same across all embedding methods, having three linear layers where the last layer returns a binding affinity score (Fig. 1b). Taking `catELMo` as an example, we first obtain a `catELMo` representation of length 1,024 for each sequence. We then feed the TCR representation to a linear layer with 2,048 neurons, followed by a Sigmoid Linear Units (SiLU) activation function [58], batch normalization [59], and 0.3 rate dropout [60]. Similarly, we feed the epitope representation to another linear layer with 2,048 neurons, followed by the same layers. The outputs of TCR and epitope layers are then concatenated (4,096 neurons) and passed into a linear layer with 1,024 neurons, followed by a SiLU activation function, batch normalization, and 0.3 rate dropout. Finally, we append the last linear layer with a neuron followed by a sigmoid activation function to obtain the binding affinity score ranging from 0 to 1. The models are trained to minimize a binary cross-entropy loss via Adam optimizer [61]. We set the batch size as 32 and the learning rate as 0.001. We stop the training if either the validation loss does not decrease for 30 consecutive epochs or it iterates over 200 epochs. Finally, we compare and report AUC scores of binding affinity prediction models of the different embedding methods.

#### 4.4.2 Epitope-specific TCR clustering

Clustering TCRs is the first and fundamental step in TCR repertoire analysis as it can potentially identify TCR clonotypes that are condition-specific [36, 37]. We perform hierarchical clustering [31] to `catELMo` and the state-of-the-art amino acid embeddings (further demonstrated in Section 4.5). We also obtain clusters from the existing TCR clustering approaches (TCRdist and GIANA). Both methods are developed on the BLOSUM62 matrix and apply nearest neighbor search to cluster TCR sequences. GIANA used the CDR3 of TCR*β* chain and V gene, while TCRdist predominantly experimented with CDR1, CDR2, and CDR3 from both TCRα and TCR*β* chains. For fair comparisons, we perform GIANA and TCRdist only on CDR3 *β* chains and with hierarchical clustering instead of the nearest neighbor search.

We first obtain different types of TCR embeddings from `catELMo` and the comparison methods. All embedding methods except BLOSUM62 yield the same size representation vectors regardless of TCR length. For BLOSUM62 embedding, we pad the sequences so that all sequences are mapped to the same size vectors (further demonstrated in Section 4.5). We then perform hierarchical clustering on TCR embeddings of each method. In detail, the clustering algorithm starts with each TCR as a cluster with size 1. It repeatedly merges the closest two clusters based on the Euclidean distance between TCR embeddings until it reaches the target number of clusters.

We compare the normalized mutual information (NMI) between the identified cluster and the ground-truth.

NMI is a harmonic mean between homogeneity and completeness. Homogeneity measures how many TCRs in a cluster bind to the same epitope, while completeness measures how many TCRs binding to the same epitope are clustered together. A higher value indicates a better clustering result. It ranges from zero to one where zero indicates no mutual information found between the identified clusters and the ground-truth clusters and one indicates a perfect correlation.

### 4.5 Comparison studies

We demonstrate how we implement existing amino acid embedding methods to compare with catELMo for the two TCR-related downstream tasks.

#### BLOSUM62

Among various typs of BLOSUM matrices, we use BLOSUM62 as it has been widely used in many TCR-related models [7, 9, 11, 17, 37, 38]. We obtain embeddings by mapping each amino acid to a vector of length 24 via BLOSUM62 matrix. Since TCRs (or epitopes) have varied lengths of the sequences, we pad each sequence using IMGT [62] method. If a TCR sequence is shorter than the predefined length 20 (or 22 for epitopes) [10], we add zero-padding to the middle of the sequence. Otherwise, we remove amino acids from the middle of the sequence until it reaches the target length. For each TCR, we flatten 20 amino acid embedding vectors of length 24 into a vector of length 480. For each epitope, we flatten 22 amino acid embedding vectors of length 24 into a vector of length 528.

#### Yang et al

We select the 3-mer model with a window size of 5 to embed TCR and epitope sequences, which is the best combination obtained from a grid search. Each 3-mer is embedded as a numeric vector of length 64. The vectors are averaged to represent a whole sequence, resulting in a vector of length 64.

#### SeqVec and ProtBert

We embed each amino acid as a numeric vector of length 1,024. The vectors are element-wisely averaged to represent a whole sequence, resulting in a vector of length 1,024.

#### TCRBert

We embed each amino acid as a numeric vector of length 768. The vectors are element-wisely averaged to represent a whole sequence with a vector of length 768.

### 4.6 Ablation studies

We provide details of our experimental design and ablation studies.

#### 4.6.1 Depth of catELMo

We investigate the effect of various depths of `catELMo` on TCR-epitope binding affinity prediction performance. We compare `catELMo` with different numbers of BiLSTM layers, specifically `catELMo`-Shallow, `catELMo`, and `catELMo`-Deep with 2, 4 and 8 layers respectively. Other hyperparameters and the training strategy remained the same as described in Section 4.3. For each amino acid residue, we average the output vectors of CharCNN and four (or two, eight) BiLSTM, resulting in a numerical vector of length 1,024. We then element-wisely average over all amino acids’ representations to represent a whole sequence, resulting in a numerical vector of length 1,024. Embeddings from various depths are used to train binding affinity prediction models, resulting in three sets of downstream models. All settings of the downstream models remain the same as described in Section 4.4.1. The downstream models’ prediction performance is compared to investigate the optimal depth of `catELMo`.

#### 4.6.2 Neural architecture of catELMo

We compare `catELMo` with BERT-based amino acid embedding models using another context-aware architecture, Transformer, which has shown outstanding performance in natural language processing tasks. We train different sizes of BERT, a widely used Transformer-based model, for amino acid embedding, named BERT-Base-TCR, BERT-Tiny-TCR, and BERT-Large-TCR. Each model has 2, 12, and 30 Transformer layers and returns 768, 768, and 1024 sizes of embeddings for each amino acid token. Their objectives, however, only consist of the masked language prediction and do not include the next sentence prediction. For each TCR sequence, 15% of amino acid tokens are masked out and the model is trained to recover the masked tokens based on the remaining ones. The models are trained on the same training set as `catELMo` for 10 epochs. Other parameter settings are the same as TCRBert, which is included as one of the comparison models. All other settings remain the same as described in Section 4.4.1. TCRBert and BERT-Base-TCR share the same architecture, whereas TCRBert is trained on fewer training samples (PIRD). The embedding of a whole TCR sequence is obtained by average pooling over all amino acid representations. Embeddings from each model are used to train binding affinity prediction models, resulting in three sets of downstream models. The prediction performance of the downstream prediction models is compared to evaluate the architecture of `catELMo`.

#### 4.6.3 Size of downstream data

We investigate how much downstream data `catELMo` can save in training a binding affinity prediction model while achieving the same performance with a model trained on a full size of data. We train the same model on different portion of `catELMo` embedding dataset. In detail, we randomly select k% of binding and k% of non-binding TCR-epitope pairs from training (and validation) data (k = 1, 2, ·· ·, 10, 20, ·· ·, 100), obtain `catELMo` embeddings for those, and feed them to train TCR-epitope binding affinity prediction models. Note that the TCR-epitope binding affinity prediction models in this experiment differ only in the number of training and validation pairs, meaning that the same testing set is used for different ks. We run ten times of experiments for each k and report their average and standard deviation of AUC, recall, precision, and F1 scores. We compare their performance to those trained on a full size of the other embedding datasets. For a more detailed investigation, we also perform the same experiment on BLOSUM62 embeddings and compare it with ours.

## Data availability

Datasets used in downstream tasks and embedding models are described in detail in Section 4.1. Those datasets have been combined and summarized by us and are accessible at at https://github.com/Lee-CBG/catELMo/tree/main/datasets.

## Code availability

The implementation of catELMo as well as all of our trained embedding models are publicly available at https://github.com/Lee-CBG/catELMo.

## Authors’ contributions

HL conceived the idea for the overall project. PZ, SB and HL designed the method and wrote the manuscript. PZ carried out the experiments. MC collected data and participated in project discussions.

## Ethics declarations

The authors declare no competing interests.

