## Supplementary Materials for "Context-Aware Amino Acid Embedding Advances Analysis of TCR-Epitope Interactions"

February 28, 2024

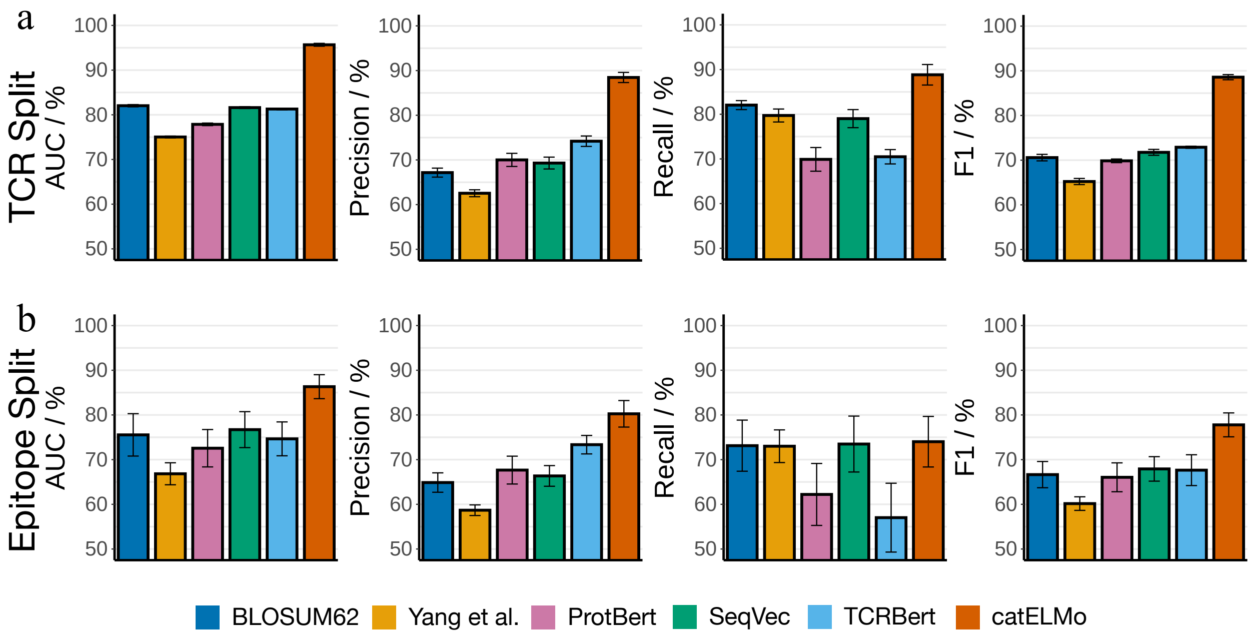


Supplementary Figure 1. **Prediction performance of TCR-epitope binding affinity prediction models trained on different embedding methods**. The average and standard deviation (error bar) of 10 trials are reported. **a**) AUC, precision, recall, and F1 of TCR split, and **b**) epitope split.

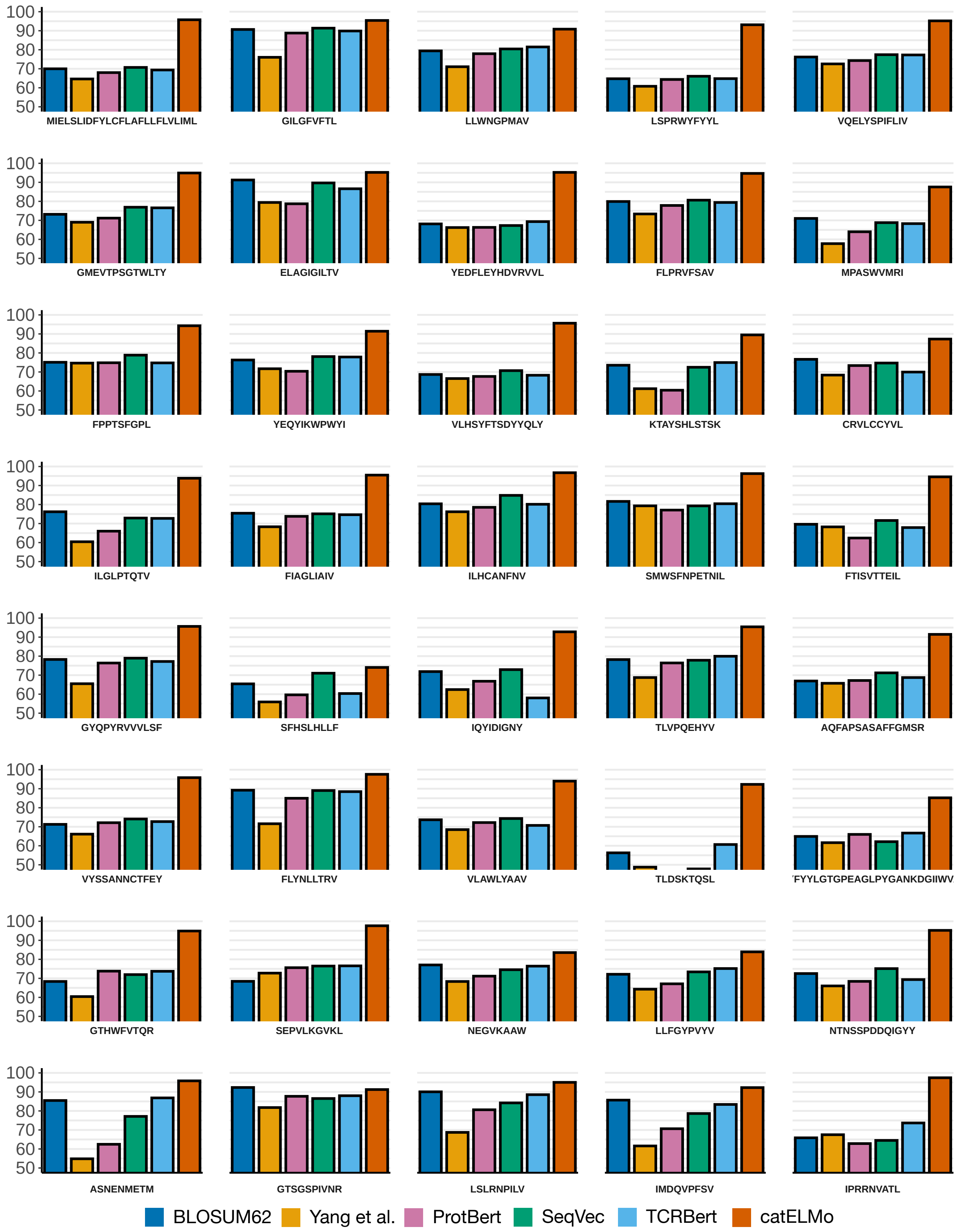


Supplementary Figure 2. **AUCs of the top 40 frequent out of sample epitopes in the binding affinity prediction task**.

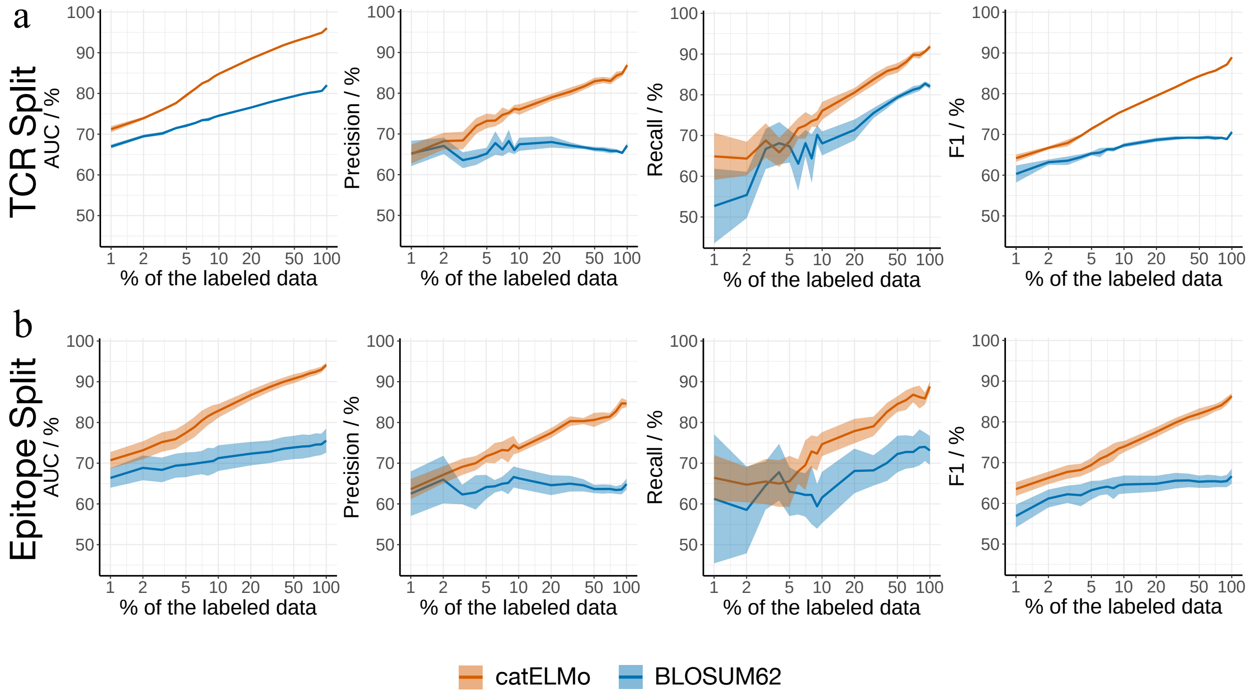


Supplementary Figure 3. **AUCs of TCR-epitope binding affinity prediction models trained on different portions of downstream datasets.**. The average and interval of 95% confidence (error band) of 10 trials are reported. **a**) AUC, precision, recall, and F1 of TCR split, and **b**) epitope split.

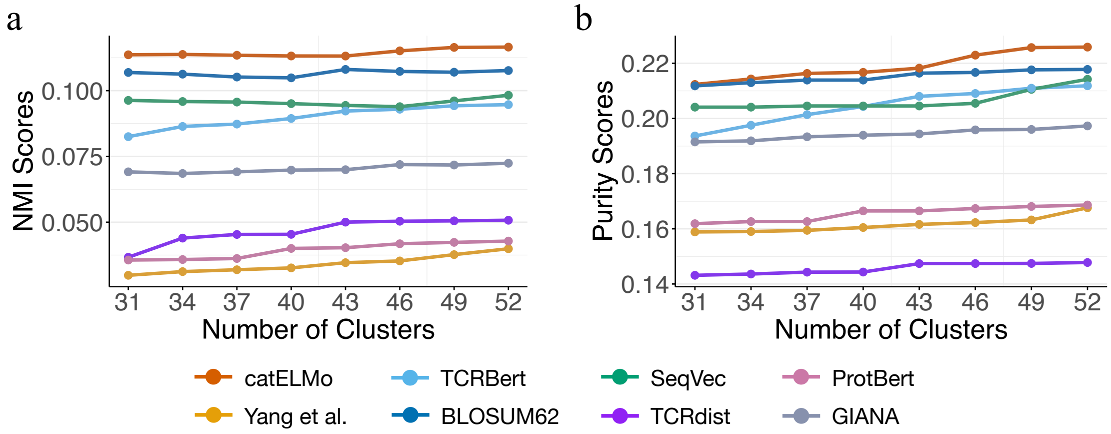


Supplementary Figure 4. **TCR clustering performance for the top 34 abundant epitopes representing 70.55% of TCRs in our collected databases.** Hierarchical clustering was employed on embeddings computed by each method. `catELMo` consistently outperformed other methods in **a**) NMI scores and **b**) cluster purity.

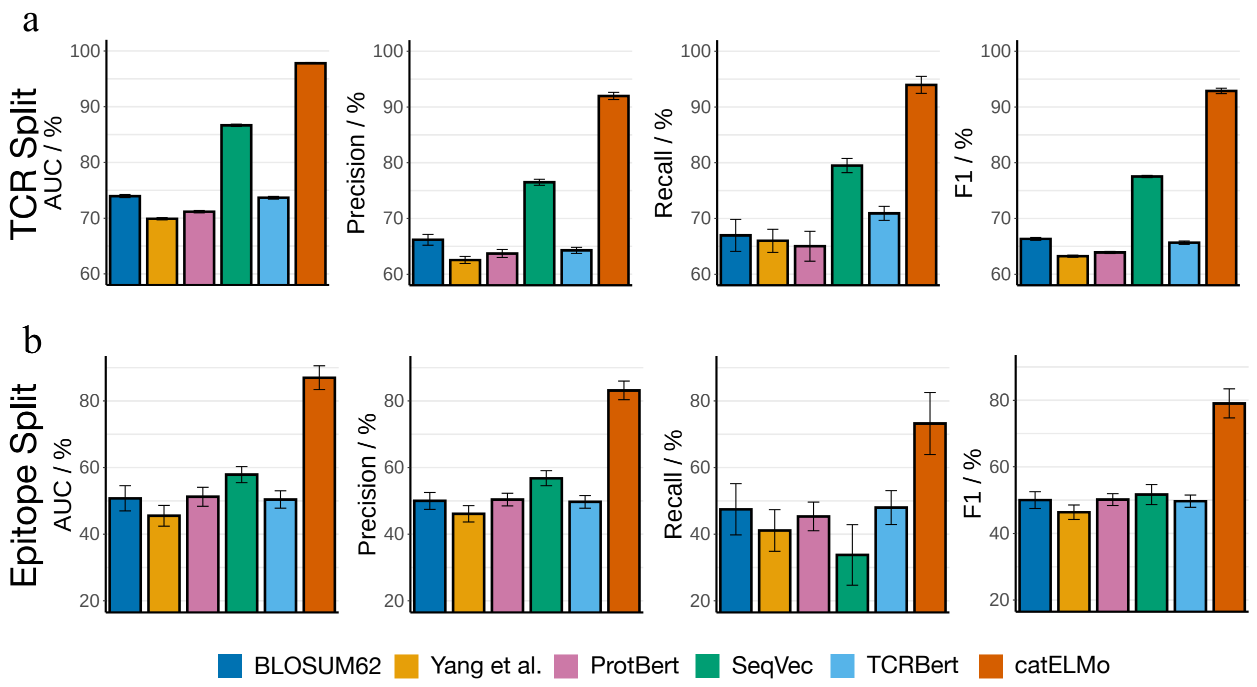


Supplementary Figure 5. **Prediction performance of TCR-epitope binding affinity prediction models trained on different embeddings where negative samples are generated by random shuffling.** The average and standard deviation (error bar) of 10 trials are reported. **a**) AUC, precision, recall, and F1 of TCR split and **b**) epitope split.

Supplementary Table 1. **AUC scores for top 10 frequent epitope types (pathogens) in the testing set of epitope split.** The highest AUC scores across six embedding models are highlighted in **bold**.

| Pathogens | Number of TCRs | BLOSUM62 | Yang et al. | ProtBert | SeqVec | TCRBert | catELMo |
|---|---|---|---|---|---|---|---|
| SARS-CoV-2 | 38606 | 70.32 | 65.25 | 68.16 | 71.25 | 69.85 | **95.1** |
| Influenza | 10802 | 90.66 | 76.03 | 88.77 | 91.36 | 89.86 | **95.4** |
| Yellow fever virus | 4716 | 79.4 | 71.07 | 77.92 | 80.42 | 81.47 | **90.83** |
| Human coronavirus (strain SARS) | 4646 | 73.78 | 66.32 | 70.01 | 74.58 | 72.15 | **92.36** |
| Melanoma | 2090 | 91.05 | 78.76 | 78.43 | 89.34 | 86.54 | **94.89** |
| SARS coronavirus Tor2 | 2058 | 78.82 | 72.57 | 76.96 | 79.73 | 78.04 | **94.51** |
| Cytomegalovirus (CMV) | 1104 | 76.49 | 68.87 | 72.46 | 73.58 | 70.76 | **86.19** |
| Hepatitis B virus (HBV) | 954 | 73.5 | 61.3 | 60.48 | 72.46 | 74.8 | **89.33** |
| Neoantigen | 850 | 80 | 69.01 | 76.26 | 80.94 | 79.79 | **89.8** |
| HTLV-1 | 370 | 65.43 | 55.93 | 59.65 | 71.03 | 60.34 | **74.05** |

Supplementary Table 2. **TCR-epitope binding affinity prediction performance of TCR split where negative samples are generated by random shuffling.** Average and standard deviation of 10 trials are reported. P-values are from two-sample t-tests between `catELMo` and the second best method (underlined).

| | AUC (%) | Precision (%) | Recall (%) | F1 (%) |
|---|---|---|---|---|
| BLOSUM62 | 73.95 ± 0.28 | 66.18 ± 0.96 | 66.97 ± 2.86 | 66.34 ± 0.24 |
| Yang et al. | 69.89 ± 0.19 | 62.54 ± 0.65 | 66.00 ± 2.08 | 63.24 ± 0.20 |
| ProtBert | 71.16 ± 0.20 | 63.69 ± 0.72 | 65.03 ± 2.68 | 63.90 ± 0.21 |
| SeqVec | <u>86.66 ± 0.22</u> | <u>76.50 ± 0.55</u> | <u>79.48 ± 1.27</u> | <u>77.52 ± 0.21</u> |
| TCRBert | 73.67 ± 0.24 | 64.29 ± 0.56 | 70.92 ± 1.26 | 65.65 ± 0.31 |
| `catELMo` (ours) | **97.80 ± 0.13** | **91.98 ± 0.65** | **93.97 ± 1.53** | **92.88 ± 0.49** |
| p-value | $1.04 \times 10^{-18}$ | $8.00 \times 10^{-14}$ | $1.77 \times 10^{-11}$ | $1.82 \times 10^{-22}$ |

Supplementary Table 3. **TCR-epitope binding affinity prediction performance of epitope split where negative samples are generated by random shuffling.** Average and standard deviation of 10 trials are reported. P-values are from two-sample t-tests between `catELMo` and the second best method (underlined).

| | AUC (%) | Precision (%) | Recall (%) | F1 (%) |
|---|---|---|---|---|
| BLOSUM62 | 50.76 ± 3.79 | 50.01 ± 2.54 | 47.47 ± 7.70 | 50.02 ± 2.49 |
| Yang et al. | 45.54 ± 3.13 | 46.11 ± 2.46 | 41.12 ± 6.25 | 46.38 ± 2.15 |
| ProtBert | 51.24 ± 2.85 | 50.39 ± 1.94 | 45.32 ± 4.31 | 50.18 ± 1.76 |
| SeqVec | <u>57.89 ± 2.42</u> | <u>56.79 ± 2.26</u> | 33.77 ± 9.09 | <u>51.69 ± 3.00</u> |
| TCRBert | 50.41 ± 2.59 | 49.71 ± 1.91 | <u>48.00 ± 5.08</u> | 49.69 ± 1.83 |
| `catELMo` (ours) | **86.95 ± 3.58** | **83.16 ± 2.82** | **73.24 ± 9.31** | **79.03 ± 4.36** |
| p-value | $6.26 \times 10^{-14}$ | $1.17 \times 10^{-13}$ | $2.85 \times 10^{-6}$ | $8.65 \times 10^{-12}$ |

Supplementary Table 4. **AUCs of TCR-epitope binding affinity prediction model trained on different portions of downstream `catELMo` embeddings where negative samples are generated by random shuffling.** The average and standard deviation of 10 trials are reported.

| Percentile of Training Samples Used | TCR Split (%) | Epitope Split (%) |
|---|---|---|
| 1% | 68.17 ± 0.60 | 51.14 ± 5.76 |
| 2% | 75.11 ± 0.69 | 55.21 ± 6.09 |
| 3% | 78.92 ± 0.63 | 58.06 ± 6.82 |
| 4% | 81.99 ± 0.52 | 61.55 ± 6.18 |
| 5% | 83.51 ± 0.73 | 63.34 ± 4.72 |
| 6% | 85.19 ± 0.65 | 65.57 ± 5.91 |
| 7% | 86.34 ± 0.55 | 67.38 ± 4.31 |
| 8% | 87.38 ± 0.55 | 69.03 ± 5.57 |
| 9% | 88.16 ± 0.46 | 70.14 ± 4.99 |
| 10% | 88.56 ± 0.48 | 71.64 ± 4.09 |

Supplementary Table 5. **AUCs of TCR-epitope binding affinity prediction models with BLOSUM62 to embed epitope sequences**. The average and standard deviation of 10 trials are reported.

| TCR Embedding | Epitope Embedding | TCR Split (%) | Epitope Split (%) |
|---|---|---|---|
| BLOSUM62 | BLOSUM62 | $82.03 \pm 0.25$ | $75.54 \pm 4.74$ |
| Yang et al. | BLOSUM62 | $74.85 \pm 0.06$ | $69.12 \pm 0.24$ |
| ProtBert | BLOSUM62 | $78.02 \pm 0.07$ | $72.1 \pm 0.48$ |
| SeqVec | BLOSUM62 | $81.67 \pm 0.19$ | $76.3 \pm 0.22$ |
| TCRBert | BLOSUM62 | $81.07 \pm 0.12$ | $74.41 \pm 0.42$ |
| `catELMo` | BLOSUM62 | $96.30 \pm 0.15$ | $94.46 \pm 1.08$ |

Supplementary Table 6. **AUCs of TCR-epitope binding affinity prediction models trained on `catELMo` TCR embeddings and random-initialized epitope embeddings**. The average and standard deviation of 10 trials are reported.

| TCR Embedding | Epitope Embedding | TCR Split (%) | Epitope Split (%) |
|---|---|---|---|
| `catELMo` | `catELMo` | $96.04 \pm 0.12$ | $94.10 \pm 0.90$ |
| `catELMo` | Randomization | $96.19 \pm 0.09$ | $94.61 \pm 0.11$ |

Supplementary Table 7. **AUCs of TCR-epitope binding affinity prediction models trained on `catELMo` and BLOSUM62 embeddings**. The average and standard deviation of 10 trials are reported.

| TCR Embedding | Epitope Embedding | TCR Split (%) | Epitope Split (%) |
|---|---|---|---|
| `catELMo` | `catELMo` | $96.04 \pm 0.12$ | $94.10 \pm 0.90$ |
| `catELMo` | BLOSUM62 | $96.30 \pm 0.15$ | $94.46 \pm 1.08$ |
| BLOSUM62 | `catELMo` | $82.16 \pm 0.17$ | $75.30 \pm 4.49$ |
| BLOSUM62 | BLOSUM62 | $82.03 \pm 0.25$ | $75.54 \pm 4.74$ |
